# Drought tolerance of *Hakea* species (Proteaceae) from a range of biomes and life-histories predicted by climatic niche

**DOI:** 10.1101/2020.09.01.276931

**Authors:** Osazee O. Oyanoghafo, Corey O’ Brien, Brendan Choat, David Tissue, Paul D. Rymer

## Abstract

Extreme drought conditions across the globe are impacting biodiversity with serious implications for the persistence of native species. However, quantitative data on drought tolerance is not available for diverse flora to inform conservation management. We quantified physiological drought tolerance in the diverse Hakea genus (Proteaceae) to test predictions based on climatic-origin, life history and functional traits. We sampled terminal branches of replicate plants of 16 species in a common garden. Xylem cavitation was induced in branches under varying water potential (tension) in a centrifuge and the tension generating 50% loss of conductivity (stem P50) was characterized as a metric for drought tolerance. The same branches were used to estimate plant functional traits, including wood density, specific leaf area, and Huber value (sap flow area to leaf area ratio). There was significant variation in stem P50 among species, which was negatively associated with the species climate-origin (rainfall and aridity). Drought tolerance did not differ among life histories; however, a drought avoidance strategy with terete leaf form and greater Huber value may be important for species to colonize and persist in the arid biome. Our findings will contribute to future prediction of species vulnerability to drought and adaptive management under climate change.

## Introduction

The impacts of drought on diverse biomes across the globe are substantial, with prolonged drought resulting in forest dieback and plant mortality, changes in species distribution, local extinction and decline in ecosystem function and resilience (Allen et al., 2010; Goulden and Bales, 2019; Powers et al., 2020). Predicting the impacts of drought on biomes and plant lineages remains a challenging task for scientists, as most predictions relying on species distribution models (SDM) and climatic niche data lack the species physiological tolerance (Fitzpatrick et al., 2008; McDowell et al., 2008; Razgour et al., 2019; Urban, 2015). Hence, quantifying species physiological thresholds is key to understanding how plants will cope with extreme climatic-induced events such as drought in the future (Allen et al., 2010).

One promising strategy to quantify physiological tolerance to drought is by characterizing hydraulic traits in relation to water limitation (Choat et al., 2012; Martin-StPaul et al., 2017). This is particularly important, as studies have shown that most flowering plants (angiosperms) function close to their hydraulic safety margin (minimum xylem pressure experienced in the field - water potential causing 50% loss of conductivity (P_50_)), and are vulnerable to climate change (Choat et al., 2012). Under prolonged drought conditions, stomatal closure is unable to prevent the continuous decline of the xylem pressure, leading to cavitation, a phase change from liquid water to gas, and the formation of gas emboli (Choat et al., 2012). This results in loss of xylem hydraulic conductivity, and in severe cases, hydraulic failure and subsequent mortality (Anderegg et al., 2016; Brodribb and Cochard, 2009; Cochard, 2014; McDowell et al., 2008; Pockman et al., 1995; Urli et al., 2013). A large body of evidence has shown species drought tolerance is quantitatively linked with resistance to cavitation in woody species (Adams et al., 2017; Brodribb and Cochard, 2009; Choat et al., 2012; Kursar et al., 2009; McDowell et al., 2008; Pockman et al., 1995). The xylem tensions associated with irreversible damage (hydraulic failure) are approximated by P_50_ in gymnosperms and by P_88_ (i.e. water potential at 88% loss of conductivity) in angiosperms (Anderegg et al., 2016; Urli et al., 2013), possibly reflecting structural and functional differences in water transport systems (Choat et al. 2018).

Plants have adapted to water deficit through a wide range of life history and functional traits, with underlying anatomical and physiological mechanisms enabling them to colonise and persist in variable climate. P_50_ is known to be correlated with life history (Pratt et al., 2007), structure and function (Brodribb and Holbrook, 2004; Jacobsen et al., 2007), and species climate range (Bourne et al., 2017). For instance, in drier climates species tend to have higher wood density, which provides greater resistance to xylem conduit implosion under high xylem tensions and is strongly correlated with P_50_ (Barotto et al., 2018; Hacke et al., 2001; Jacobsen et al., 2005). Huber value (HV: ratio of sapwood area to leaf area) is observed to be negatively related to site water availability and P_50_, such that species in drier climates have higher HV and more negative P_50_ than species in wetter climates (Gotsch et al., 2010; Markesteijn et al., 2011). Leaf size has been observed to be negatively related to drought tolerance (P_50_) such that species with small leaves tend to be more cavitation resistant (Markesteijn et al., 2011; Schreiber et al., 2016). Studies have quantified and explored species vulnerability to climate-induced drought (P_50_) in relation to functional traits across biomes (Blackman et al., 2017, 2014; Bourne et al., 2017; Larter et al., 2017; Li et al., 2019, 2018; Lucani et al., 2019; Martorell et al., 2014; Nardini and Luglio, 2014; Pita et al., 2003). However, our knowledge on the drought tolerance (P_50_) of diverse related species in relation to the interactive effects of functional and life-history traits on species survival across biomes remain limited.

The *Hakea* genus is an ideal candidate for exploring variation in physiological drought tolerance across contrasting biomes. This is because Hakea is one of the two genera (the other being *Grevillea*) within the ancient Gondwana plant family Proteaceae, that have successfully transitioned into the arid biome in the Australian continent. Hakea also display a wide variation in functional and life history traits within and among biomes. For instance, some species re-sprout either from root suckers, epicormic or lignotuber buds after disturbance such as fire and drought (e.g. *H. purpurea, H. drupacea* and *H. bakeriana*), while other *Hakea* species must rely on seed production (e.g. *H. sericea*) (Clarke et al., 2013; Groom and Lamont, 1996; Weston, 1995). Leaf morphology varies greatly among species, with broad-leaved (e.g. *H. dactyloides*, *H. cristata, H. bucculenta*) and terete leaved species (e.g. *H. leucoptera, H. tephrospermum, H. sericea*) (Groom and Lamont, 1996). The differences in functional traits and life history forms among the genus could influence species response to stress conditions (Groom and Lamont, 1996; Zeppel et al., 2015). Studies have shown that resprouting species tend to allocate more biomass to roots than shoots (Moreira et al., 2012; Pausas et al., 2016), as well as exhibiting lower rates of photosynthesis, hydraulic conductivity, and transpiration (Hernández et al., 2011; Vilagrosa et al., 2014). Leaf shape influences species exposure to drought, as such within warmer and drier sites, species tend to be needle-leaved (Groom and Lamont, 1996; Wright et al., 2017). Resprouting capacity and needle-leaves support a drought avoidance strategy, and as such may have variable drought tolerance (Groom and Lamont, 1996; Vilagrosa et al., 2014; Zeppel et al., 2015).

In this study we aimed to determine the drought tolerance of *Hakea* species that differ in life history and climatic niches to investigate what attributes are predictive of aridity. Firstly, we hypothesized that there will be significant variation in drought tolerance between species, and that this would be predicted by different life-histories. Specifically, non-resprouting species will have higher P_50_ than resprouting species, and that needle-leaved species will have higher P_50_ than broad-leaved species (Groom and Lamont, 1996; Hernández et al., 2011; Vilagrosa et al., 2014). Secondly, we hypothesized that drought tolerance (P_50_) will be predicted by species climate such that species in drier climates will have higher P_50_ than species in wetter climates (Bourne et al., 2017; Larter et al., 2017; Trueba et al., 2017). Thirdly, drought tolerance (P_50_) will be positively correlated to Huber Value (HV) and wood density (WD), and negatively correlated to specific leaf area (SLA). This study will therefore provide empirical evidence on species drought tolerance (P_50_) to inform conservation management of diverse native flora.

## Materials and Methods

### Experimental design and Species selection

All samples were collected from the same site, the Australian Botanic Garden (ABG), Mount Annan, NSW, Australia (GPS location: Lat. −34.0703, Log. 150.7668, average annual rainfall of 759 mm (2007-2016)) and were well-watered via irrigation systems (simulating a common garden design). Comparing multiple species from the same *Hakea* genus in a common garden minimizes environmental effects and allows quantification of genetically determined trait variation. Using this approach, we examined the variation in functional and hydraulic traits of a diverse array of species sampled from across the *Hakea* phylogeny (Cardillo et al. 2017). A total of 16 species were selected to represent a wide range of vegetation type, biome, climate, and life histories (Table 1). Species occurrence records were downloaded from the Australian Living Atlas (ALA) (https://www.ala.org.au/, 2019). Vegetation type was defined according to the World Wildlife Fund (WWF) as abbreviated by Cardillo et al. (2017); Arid (Deserts and Xeric Shrublands), Mediterranean (Mediterranean Forests, Woodlands and Scrub), Forest (Temperate Broadleaf and Mixed Forests), Grasslands (Temperate Grasslands, Savannas, and Shrublands). The vegetation harboring greater than 50% of the species occurrence records was assigned as its vegetation type. Biome was defined based on the aridity index (UNEP, 1997) as broadly humid and arid (aridity index > 0.5, < 0.5, respectively). The climate summary details for each species distribution was obtained from The Atlas of Living Australia using R v3.6.3 (RCoreTeam, 2020).

**Table 1:** Hakea species investigated showing the life-histories (resprouting ability, leaf form), dominant vegetation type (WWF), biome, mean annual temperature (MAT, °C), mean annual precipitation (MAP, mm), and mean aridity index (AI).

| Species | Resprouting<br>ability | Leaf form | Vegetation | Biome | MAT | MAP | AI |
| --- | --- | --- | --- | --- | --- | --- | --- |
| *Hakea archaeoides* | Resprouter | broadleaved | Forest | Humid | 16.82 | 1404 | 0.96 |
| *Hakea bucculenta* | Non-sprouter | broadleaved | Mediterranean | Arid | 19.56 | 430.9 | 0.16 |
| *Hakea cristata* | Resprouter | broadleaved | Mediterranean | Arid | 16.66 | 826.5 | 0.41 |
| *Hakea dactyloides* | Non-sprouter | broadleaved | Forest | Humid | 15.24 | 1033.6 | 0.76 |
| *Hakea eyreana* | Resprouter | Terete | Arid | Arid | 22.12 | 217.6 | 0.08 |
| *Hakea grammatophylla* | Resprouter | broadleaved | Arid | Arid | 20.53 | 330.9 | 0.13 |
| *Hakea ivoryi* | Resprouter | Terete | Grassland | Arid | 20.38 | 361.9 | 0.14 |
| *Hakea leucoptera* | Resprouter | Terete | Arid | Arid | 19.41 | 294.9 | 0.13 |
| *Hakea microcarpa* | Non-sprouter | Terete | Forest | Humid | 11.58 | 1015.4 | 0.85 |
| *Hakea tephrospermum* | Resprouter | Terete | Grassland | Arid | 17.44 | 424.2 | 0.2 |
| *Hakea bakeriana* | Resprouter | Terete | Forest | Humid | 16.64 | 1123.7 | 0.73 |
| *Hakea francisiana* | Non-sprouter | broadleaved | Mediterranean | Arid | 18.09 | 303.3 | 0.13 |
| *Hakea gibbosa* | Non-sprouter | Terete | Forest | Humid | 16.38 | 1202.1 | 0.82 |
| *Hakea macraeana* | Non-sprouter | Terete | Forest | Humid | 13.61 | 970.2 | 0.8 |
| *Hakea salicifolia* | Non-sprouter | broadleaved | Forest | Humid | 15.39 | 1191.8 | 0.81 |
| *Hakea trifurcata* | Non-sprouter | Terete | Mediterranean | Arid | 17.03 | 588.4 | 0.33 |

### Sampling of Plant material

Three individuals for each species were sampled from the AGB. Terminal full sunlight, north-facing branch that were *ca.* 90 cm long were sampled and placed into a black plastic bag with wet tissue paper and transported immediately to the laboratory (<90 minutes). Samples were stored in a cold room at 4^°^C until they were processed (within 10 days). A standardized 50 cm branch was cut under water from the terminal end of the collected samples, from which the bottom 10 cm was excised, barked removed to estimate the sap flow area and then oven dried to obtain the wood density (WD: oven dry mass/volume). All leaves were removed from the remaining 40 cm branch and leaf area measured using the Li Cor 3100 leaf area meter. Leaf material was oven dried at 70 C for 48 h prior to obtaining the dry mass. Specific leaf area (SLA, mm^2^mg^−1^) was obtained by dividing the total leaf area by the leaf oven dry mass. The ratio of the sapwood area to leaf area was described as the Huber value (HV).

### Determination of drought tolerance

Drought tolerance was determined by vulnerability to xylem cavitation (P_50_) using the centrifuge method to induce cavitation in the xylem (Cochard et al., 2013, 2005). This advanced centrifuge technique creates centrifugal force that generates tension in the branch xylem vessels to induce cavitation in branch segment, thereby allowing measurement of xylem percentage loss of conductivity at set points of tension. Straight stems, 27 cm in length and with 6 mm basal diameter, were sampled and cut under water from the remaining 40 cm-long branch segments, placed on the custom-built rotor and spun at different velocities. To control for the artefact associated with the centrifuge, initial measurements were obtained at lower pressures (−0.5 MPa = 2378 rpm) that did not induce cavitation (López et al., 2019). The percent loss of conductance (PLC) at negative xylem pressure (tension) was automatically recorded through a step-wise increase (1000 rpm each) at *ca.* 2 min stabilization time (Zhang et al., 2017) until 90-95% loss of conductivity was attained. At each new xylem pressure (tension), hydraulic conductance (Kh) was measured from 30 repeated measures. The PLC was computed as PLC = 100 × (1 − Kh/Kmax). The dependence of PLC on xylem pressure was used to generate vulnerability curves for each species and 50 % loss of conductance (P_50_) were obtained from slope of the curve using the *fitplc* R package (Duursma & Choat, 2017).

### Statistical Analysis

Stem P_50_ difference between species were tested using a linear model (*lm*), while differences between biome, life histories traits (resprouting ability and leaf forms), as well as interactions, were determined using a linear mixed effect model (*lme4*) R package (Bates et al., 2015) with species as a random variable. Residuals of models were inspected; appropriate transformations were conducted and extreme outliers were removed where necessary. ANOVA for mixed effects models was undertaken using Kenward Roger degrees of freedom approximation. Linear mixed effects model with species as random effect was used to explore predictors of cavitation resistance (P_50_). Posthoc Tukey tests were undertaken using the *emmeans* R package (Lenth, 2020) to determine which species and life histories are significantly different.

## Results

### Variation in stem P_50_ between species, biome, and vegetation

There were significant differences in stem P_50_ between *Hakea* species (P < 0.001, R^2^ = 0.98, Table 2; Fig S5 supporting information). There was a continuous variation in P_50_ across the 16 species sampled and P_50_ varied 1.9-fold among species from −;4.27 MPa (minimum P_50_, *H. archaeoides* to −;7.99 MPa (maximum P_50_, *H. grammatophylla*) (Fig 1). Vegetation type was a significant factor in determining stem P_50_ (P = 0.03, R^2^ = 0.45; Table 2), such that species from the arid (−6.61 ± 0.42), mediterranean (−6.89 ± 0.23) and grassland vegetation (−7.15 ± 0.16) were more drought tolerant than forest species (−5.17 ± 0.17). As predicted, there were significant differences in stem P_50_ between biomes (P = 0.002, Table 2, Fig 1), such that species in arid biomes (−6.86± 0.18) were more drought tolerant than species in humid biomes (−5.17 ± 0.17, Fig 1). Biome differences explained 47% of the variation in stem P_50_. The arid biome had greater variation among species (−7.99 MPa *H. grammatophylla* to −;5.07 ± 0.06 MPa *H. eyreana*; Table 3), compared to the humid biome (−6.65 MPa *H*. *macraeana* to −;4.27 MPa *H. archaeoides*; Table 3).

**Table 2:** Analysis of variance for stem P_50_ testing for differences among species, biomes, vegetation, resprouting ability and leaf form.

| **Factors** | **F** | **Df** | **R<sup>2</sup>** | **P** |
| --- | --- | --- | --- | --- |
| Species<sup>1</sup> | 145.81 | 15 | 0.98 | 2.20E-16 |
| Biome | 14.18 | 1 | 0.47 | 0.002 |
| Vegetation | 4.31 | 3 | 0.45 | 0.028 |
| Resprouting ability | 0.17 | 1 | 0.01 | 0.688 |
| Leaf form | 0.02 | 1 | 0.00 | 0.888 |
| Biome | 20.61 | 1 |  | 0.0007 |
| Leaf form | 0.03 | 1 | 0.63 | 0.866 |
| Leaf form : Biome | 8.41 | 1 |  | 0.013 |
| Biome | 13.47 | 1 |  | 0.003 |
| Resprouting ability | 0.76 | 1 | 0.47 | 0.272 |
| Resprouting ability : Biome | 0.12 | 1 |  | 0.732 |

Each section represents a single model, single factor analysis or analysis with two factors and interaction term. <sup>1</sup> linear model (lm) for species, while all other models are include speceis as a random factor (lme).

**Fig 1:**
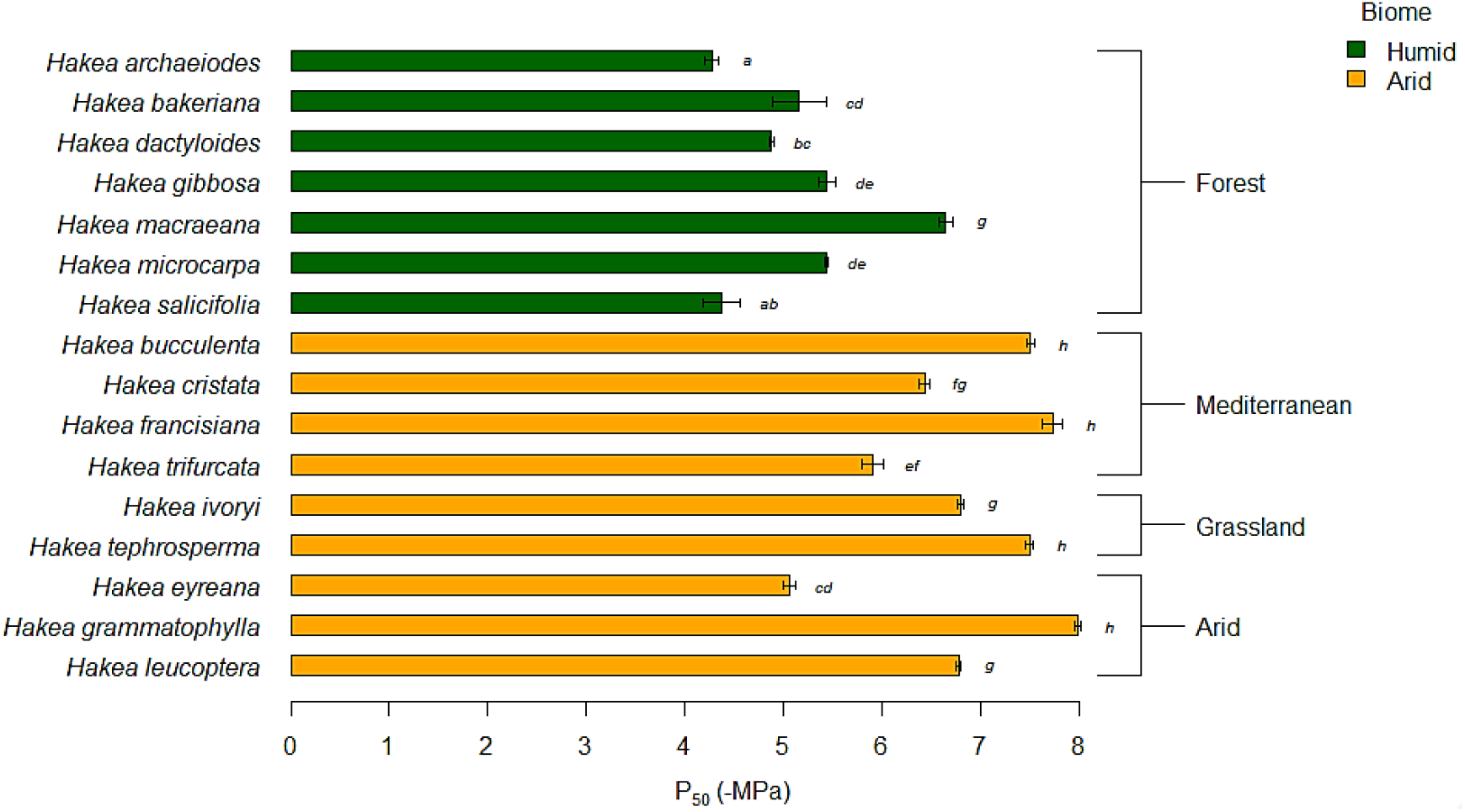
Stem P_50_ of *Hakea* species in the different biomes and vegetation types. Values presented are mean ± standard error of each species. Colours and brackets show the different biome and vegetation type, respectively. Letters denote which categories are significantly different based on a Post-hoc Tukey’s test.

**Table 3:**
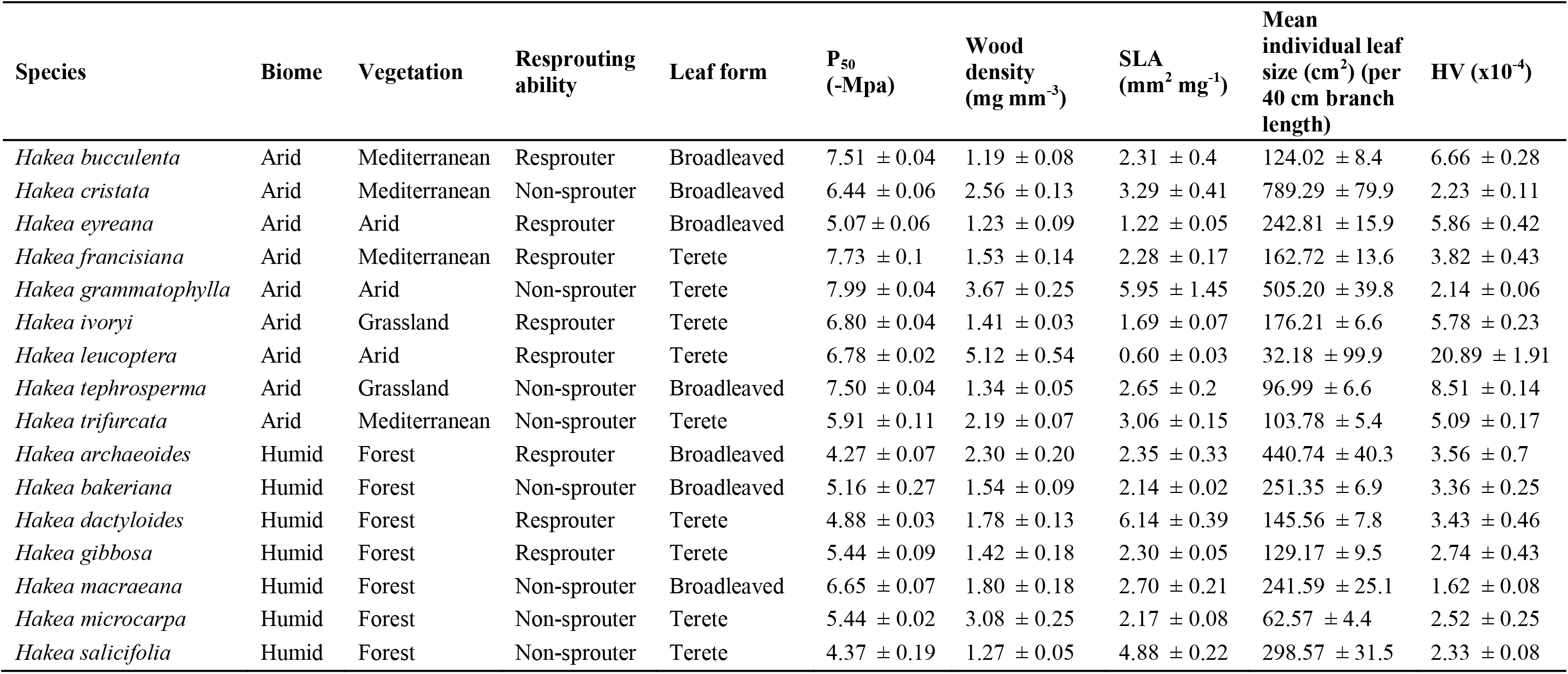
*Hakea* species biome, vegetation and life history presented alongside mean (+/− SE) cavitation resistance (stem P_50_) and functional traits.

### Cavitation resistance (stem P_50_) related to biome and life histories

There was no significant difference in P_50_ between life history types (resprouters vs non-sprouters, P = 0.688; broadleaved vs terete leaves, P = 0.888; Table 2). However, there was a significant interaction between biome and leaf form for stem P_50_ (P = 0.013; Table 2). Broadleaved species in the arid biome were significantly more drought tolerant than broad and terete leaved species in the humid biome, whilst broadleaved species in the humid biome were significantly less drought tolerant than both leaf forms in the arid biome (Fig 2). Such that drought tolerance increased from the humid biome broad to terete leaved species then to the arid biome terete to broad leaved species. No significant interaction between resprouting ability and biome was detected (Table 2), however non-sprouting and resprouting species in the arid biome were more drought tolerant than resprouters in the humid biome (Fig 2).

**Fig 2:**
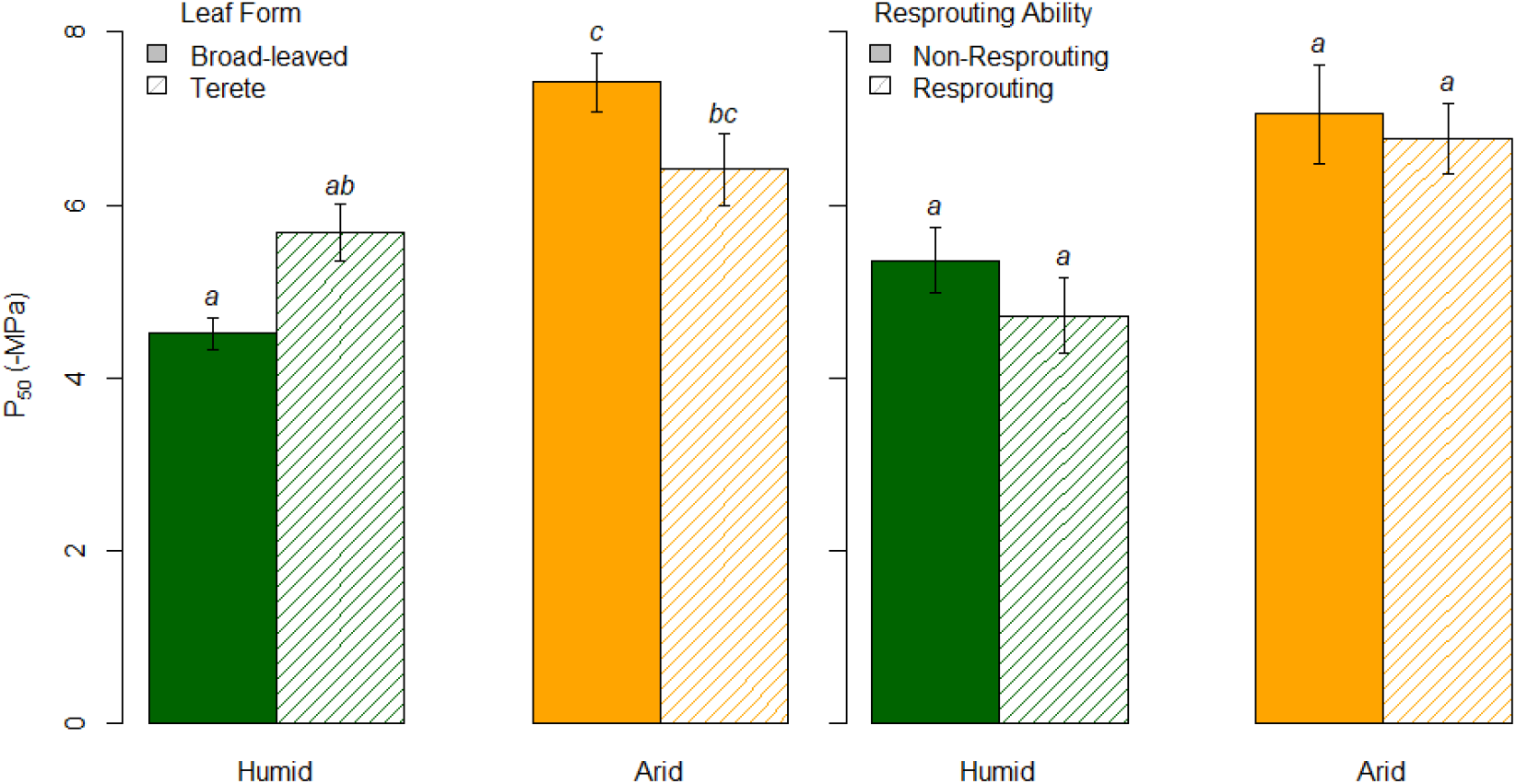
Interaction between biome and leaf form (a), and biome and resprouting ability (b) for cavitation resistance (stem P_50_ mean ± standard error). Letters denote which categories are significantly different based on a Post-hoc Tukey’s test.

### Interplay between drought tolerance, functional traits and climatic origin

P_50_ and HV were significantly correlated with species climate-origin. Drought tolerance (stem P_50_) was significantly related with rainfall (MAP R^2^ = 0.51, P-value = 0.001) and aridity (AI R^2^ = 0.49, P-value = 0.001), but unrelated with temperature (MAT R^2^ = 0.13, P-value = 0.143) (Fig 3 a, b, c). Variation in stem P_50_ was not significantly related to any functional traits; WD, SLA, leaf area and HV (Fig S4, supporting information). Similar to stem P_50_, HV was significantly related to MAP (R^2^ = 0.24, P = 0.04), AI (R^2^ = 0.22, P = 0.04) but unrelated with MAT (R^2^ = 0.156, P = 0.111) (Fig 3 d, e, f). Wood density, SLA and leaf size were not found to be significantly related to any of the climate variables.

**Fig 3:**
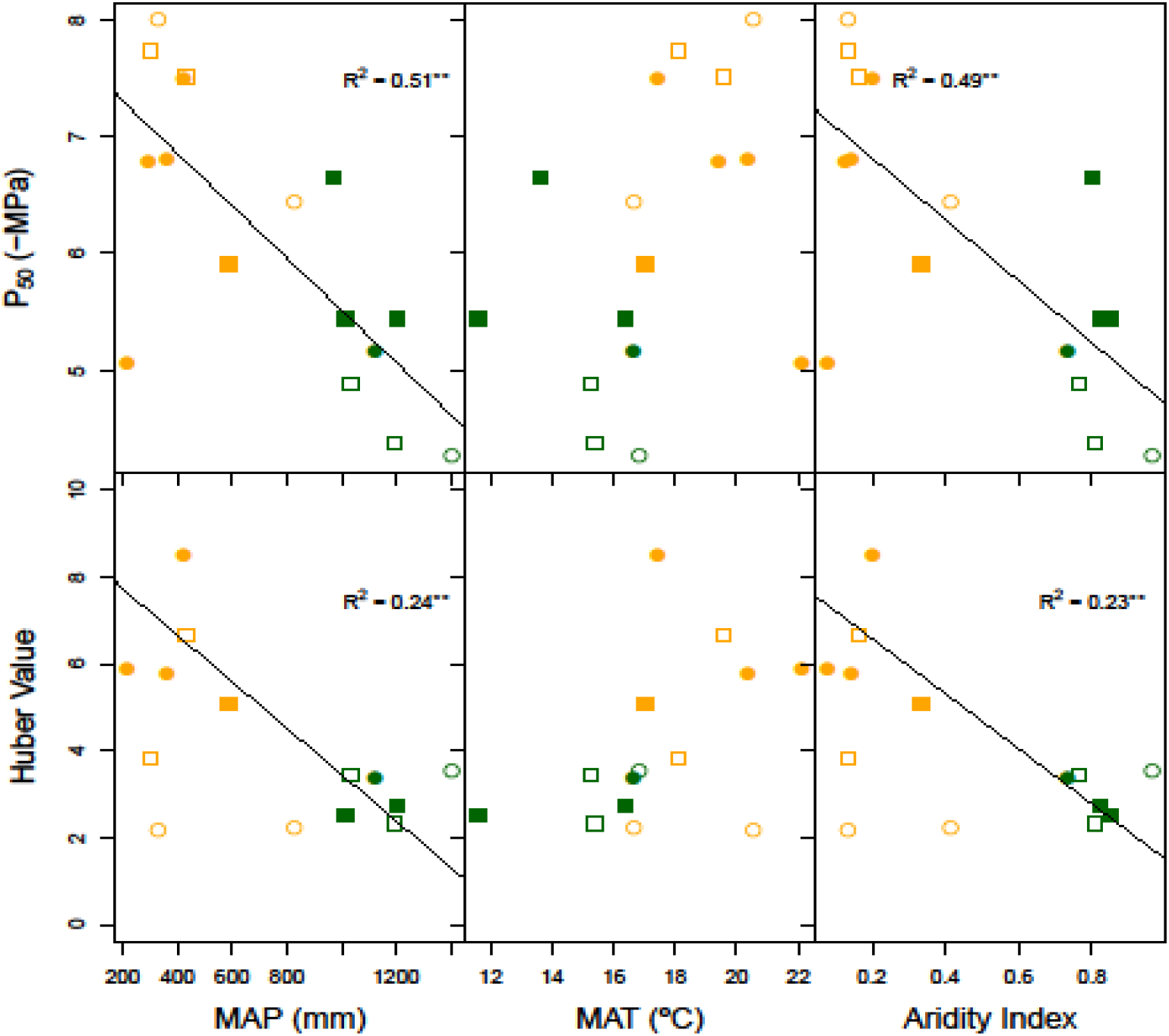
Relationship between stem water potential causing 50% loos of conductance (P_50_; panels a, b, c) and Huber Value (HV x 10^-4; panels d, e, f) with rainfall (MAP), temperature (MAT) and aridity (AI) of the species climate-origin. Values presented are mean of each species colour coded for different biomes (green = humid, orange = arid) with different shapes for life-histories (circle = resprouters, rectangle = non-resprouters) and different fill for leaf form (filled = terete, open = broadleaved). **Significance codes: ‘**’ P < 0.01, ‘*’ P < 0.05.**

## Discussion

The aim of this study was to determine the drought tolerance of a diverse array of *Hakea* species to test prediction based on climate-origin and life-history. Sampling 16 *Hakea* species representing a wide range of climatic niche and life history traits from common conditions estimated genetically determined trait variation. Our results revealed that there was significant variation in the drought tolerance of congeneric species and that biome/climate (rainfall and aridity) of origin was the key predictor of hydraulic traits (stem P_50_). Other traits contributed to drought tolerance; terete leaf form and higher sapwood area to leaf area ratio (HV) would be expected to reduce whole plant exposure and stress during periods of high evaporative demand. The study is timely given the recent devastating drought episodes experienced across the Australian continent, and in many regions throughout the world.

### Climate is a major driver of variation in cavitation resistance

Climate has often been highlighted as the key driver of species variation in hydraulic traits (Li et al., 2018). Hydraulic traits appear to be adaptive with species that have shorter and narrower vessels tending to occupy drier biomes and have lower vulnerability to cavitation (Christman et al., 2009; Larter et al., 2017; Lens et al., 2011, 2009; Pockman and Sperry, 2000; Skelton et al., 2018; Sperry et al., 2008; Wheeler et al., 2007). In this study, species in the arid biome/climate (with the potential exception of *H. eyreana*) generally had higher cavitation resistance (P_50_ below −;6.75 MPa) compared with species in the humid biome (Table 3). This was indeed expected as species within the arid biome possess traits that confer greater drought tolerance (Choat et al., 2012; Li et al., 2019, 2018; Trueba et al., 2017). To understand if drought tolerance is genetically determined hydraulic trait, we plotted P_50_ against mean annual precipitation (MAP), temperature (MAT), and aridity (AI) across the species distribution range (Fig 3). We found evidence supporting our expectation, that drought tolerance trait P_50_ is significantly related with climate (Bourne et al., 2017; Choat et al., 2012; Larter et al., 2017; Li et al., 2018; Maherali et al., 2004; Trueba et al., 2017). Our results revealed that rainfall and aridity were key drivers of the variation in species P_50_ within the diverse *Hakea* genus, such that species cavitation resistance increases with reduced rainfall of species climatic origin. Furthermore, the P_50_ trait variation measured in a common garden provides strong evidence that drought tolerance is genetically-determined and adaptive (Lamy et al., 2014; Li et al., 2018; López et al., 2016; Skelton et al., 2019). Hence, cavitation is a key factor shaping species distribution with respect to water availability (Brodribb and Hill, 1999).

### Variation in species cavitation resistance across life history traits

We observed significant species-specific variation in cavitation resistance among *Hakea* species (Table 2), demonstrating that species within the genus vary broadly in their capacity to tolerate high levels of water stress. Differences in species drought tolerance (stem P_50_) are largely attributed to the differences in the xylem structure; e.g. pit membrane porosity and thickness, and conduit size (Choat et al., 2012, 2008; Delzon et al., 2010; Li et al., 2016; Maherali et al., 2004; Sperry et al., 2006). At maturity, xylem conduits are dead with no possible acclimation to environmental change, making estimates of drought tolerance via embolism resistance very important to make reliable prediction under future climatic changes (Choat et al., 2012).

Species habitat preference and survival under disturbances (e.g. fire and drought) within the *Hakea* genus have previously been reported to be related with species life history and leaf form (Groom and Lamont, 1996). In contrast to our expectation, there were no significant differences between resprouters and non-resprouters, as well as broad and terete leaved species (Table 2). Previous studies have reported contrary findings in relation to resprouting ability and drought tolerance (Vilagrosa et al., 2014; Zeppel et al., 2015). The differences between these studies and our study may be due to the fact that Zeppel et al (2015) considered a large database of stem P_50_ with species across different genus, indicating that resprouters were more drought tolerant than non-resprouters, at least within Angiosperms. On the other hand, Vilagrosa et al. (2014) used field observation and stem hydraulic measurements focusing on 12 co-occurring woody species within the Mediterranean system found that resprouters were less drought tolerant than non-resprouters (i.e. seeders), in direct contrast to Zeppel et al. (2015). In contrast to these studies, we focused on a single genus across multiple biomes finding no difference in drought tolerance with resprouting ability. This finding is partially supported by Groom and Lamont (1996), who found resprouting ability among *Hakea* species within the Mediterranean biome of southwest Australia was not associated with aridity. The differences may be due to varying drought strategies (i.e. tolerators, avoiders) for survival among genera and biomes.

### Interactions between leaf form and biome

Arid plants show adaptations with small, terete, leaves, as seen in the distribution of Hakea species in the southwest Australian Mediterranean biome (Groom and Lamont 1996). While our findings do not support this overall pattern, we observed a significant interaction between leaf form and biome for drought tolerance. Broad-leaved species in the arid biome were significantly more drought tolerant compared to broad-leaved species in the humid biome (Fig 2). Broad leaves increase the surface area for carbon uptake (photosynthesis) and transpiration, however within warmer and drier sites, this may potentially lead to serious water loss or cavitation (Wright et al., 2017). Thus, broadleaved species within the arid biome would be more dependent on resistant xylem (higher P_50_) to prevent implosion. Our results also showed support for this (though not significant), as broadleaved species within the arid biome generally had higher P_50_ compared to terete leaved species in the same biome (Fig 2). This finding highlights the ability for different strategies to co-exist in the arid biome, and the importance of understanding trait coordination alongside climate drivers for drought adaptation.

### Avoidance of water stress as a possible strategy to persist in the arid biome

Huber value (HV, ratio of sapwood to leaf area) is a measure of carbon investment in xylem tissue per unit leaf area (Eamus et al., 2006; Gotsch et al., 2010; Pérez-Harguindeguy et al., 2013). Knowledge of the HV gives insight into the strategy species employed to survive in varying climate, as such reduced leaf area to sapwood area ratio implies avoidance strategy (Eamus et al., 2006). Species in the arid biome may employ drought avoidance strategy in addition to tolerance to persist in the arid biome (Fig 3). This was indeed true as HV was significantly related to climate (MAP, AI), such that species with higher HV tended to occupy areas with reduced rainfall, inferring greater demand for water transport (Choat et al., 2007; Gleason et al., 2012). Studies have also shown variation in drought tolerance traits within communities irrespective of the precipitation level (Maherali et al., 2004; McCulloh et al., 2019). This is true, as we observed wide variation in P_50_ within the Arid-Arid communities (or vegetation-biome, Table 3) driven by the low stem P_50_ value of *H. eyreana*. We also observed the wood density of *H.* eyreana (1.23 ± 0.09) to be about 3-fold smaller than *H. leucoptera* and 4.1-fold smaller than *H*. *grammatophylla* which also possesses a higher P_50_, suggesting that *H. eyreana* may probably employ a different strategy (e.g. drought avoidance) to survive in the arid biome. *Hakea eyreana* may have different trait coordination or trade-offs among traits along the water transport pathway under field scenarios or in response to the experimental (well-watered) condition (Brodribb et al., 2017; Li et al., 2018; McCulloh et al., 2019). Traits including life history (e.g. resprouting ability and leaf form), growth form (e.g. liana vs tree), stomatal regulation, soil water depth, and root depth may also offset the need for developing more negative stem P_50_ (Bartletta et al., 2016; Meinzer et al., 2009; Padilla and Pugnaire, 2007; Skelton et al., 2015). We found evidence that drought avoidance provided with terete leaf form and reduced leaf area to sapwood ratio (HV) may enable species with low drought tolerance (e.g. *H. eyreana*) the ability to persist in the arid biome. The plots of HV and P_50_ against aridity showing different life-histories (Fig 3) are informative in understanding different drought strategies. These traits may balance the need for carbon capture and growth with the demand for developing xylem resistant to drought (high P_50_, -MPa) for colonization and persistence in the arid biome.

### Relationship between cavitation resistance and functional traits

Surprisingly, we observed no significant relationship between P_50_ and functional traits (e.g. wood density, SLA, HV, LDMC; Fig S4 supporting information) (Hacke et al., 2001; Markesteijn et al., 2011; Schumann et al., 2019; Villagra et al., 2013). Of these functional traits, wood density has received more attention in relation to drought tolerance as greater structural investment (wood density) would prevent xylem implosion, and thus greater resistance to embolism (Hacke et al., 2001; Li et al., 2018; Markesteijn et al., 2011). However, some other studies have also reported no significant association between wood density and P_50_ (Larter et al., 2017; Trueba et al., 2017). The non-significant relationships between P_50_ and functional traits (e.g. WD and SLA) found by Trueba et al. (2017) may be because species were pooled from diverse communities and genera. Furthermore, there may be limited selective pressure for investment in structural strength since our study sites were well-watered, as studies have shown environment or site to be a determinant of wood density (Downes et al., 2006; Onoda et al., 2010; Roderick and Berry, 2002; Searson et al., 2004; Wimmer et al., 2002).

Leaf size traits were not important in pooling species apart in relation to stem P_50_, as both broad-leaved and terete leaved species are distributed within the arid and humid communities (Groom and Lamont, 1996). However, in combination with Huber value, leaf size may be important in highlighting species strategy to drought and across climate/biomes. For instance in arid biome, species with reduced leaf area to sapwood area employs avoidance strategy, while species with greater surface area within the same system will tend to prioritize the construction of xylem resistance to embolism (i.e higher stem P_50_) for survival. Interestingly, we did not observe significant relationships between hydraulic traits (HV and P_50_). However, the direction of the relationship was positive (R^2^ = 0.14, P value = 0.588) as expected (Carter and White, 2009; Markesteijn et al., 2011). The weak relationship suggests that not all species with higher P_50_ (tolerance) necessarily had higher HV (avoidance) (Fig S4, supporting information), as some species may either employ alternate strategies for survival.

## Conclusion

This study highlights climate (rainfall and aridity), rather than life history and functional traits, as the key predictor of variation in drought tolerance (stem P_50_). Rainfall for species origin was the best predictor of hydraulic trait, explaining variation in stem P_50_, which appears to be a major determinant of species distribution. This study also indicates that stem P_50_ is an adaptive trait, genetically determined, and hence reliable and robust for predicting species vulnerability to climate change. This provides support for climate as a predictor of species suitability under climate change using species distribution models. Our results show that *Hakea* species in humid biomes are more vulnerable to future droughts compared to species in arid biomes. Alternative avoidance or recovery strategies may still be important for diverse flora to colonise and persist in the arid biome. We provide evidence for avoidance via terete leaves and enhanced HV, however the role of resprouting in recovery from drought was not supported. Findings from this study will provide the scientific basis for adaptive management strategies for *Hakea*, including conservation of threatened and widespread species through translocations and assisted migration respectively.

## Acknowledgement

We appreciate the support of Benedict Lyte from The Royal Botanic Garden, Sydney for granting us excess to their living collections. Nzie Peter for support in data collection and Rosana Lopez for technical insights into the hydraulic techniques. Australian Postgraduate Award (Western Sydney University), Ecological Society of Australia-Holsworth Wildlife Research Endowment grant to O.O.O. This project has been supported by the New South Wales Government’s Department for Planning, Industry and Environment, Saving Our Species grant to P.D.R and D.T.

We appreciate the support of Benedict Lyte from The Australian Botanic Garden, Mount Annan for granting us excess to their collections and Nzie Peter for support in data collection and Rosana Lopez for technical insights into the hydraulic techniques. Australian Postgraduate Award (Western Sydney University), Ecological Society of Australia-Holsworth Wildlife Research Endowment grant to O.O.O and NSW Department for Planning, Industry and Environment, Saving Our Species grant to P.D.R and D.T.

## Supporting information

Fig S4: Relationship between drought tolerance (P_50_) and functional traits for *Hakea* species showing different biome, life-history and leaf form.

Fig S5: Hydraulic vulnerability curves for *Hakea* species showing the percentage loss of conductance against the stem water potential measured from the Cavitron.

