## Supplementary fig for "Drought tolerance of *Hakea* species (Proteaceae) from a range of biomes and life-histories predicted by climatic niche"

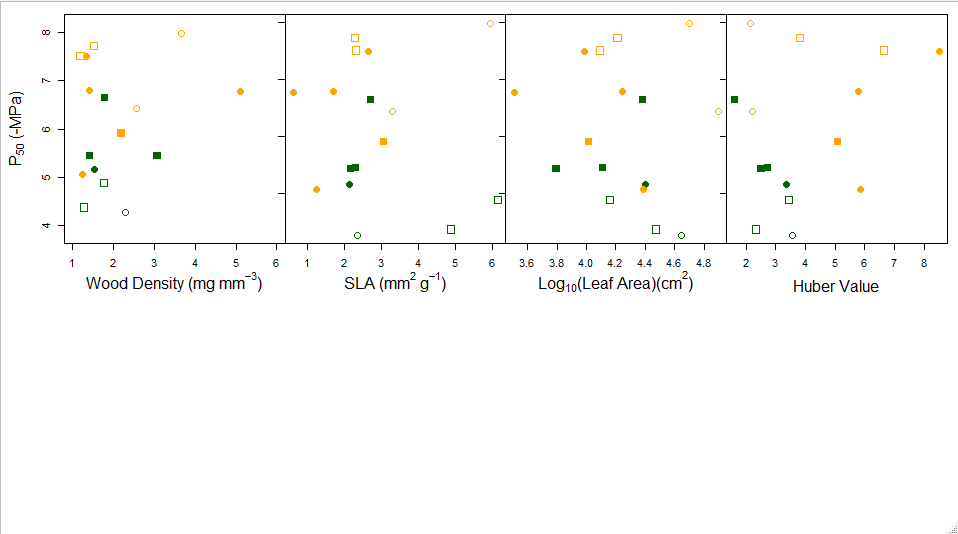

Fig S4: Relationship between drought tolerance (P_50_) and functional traits for *Hakea* species showing different biome, life-history and leaf form. Values presented are mean of each species colour coded for different biomes (green = humid, orange = arid) with different shapes for life-histories (circle = resprouters, rectangle = non-resprouters) and different fill for leaf form (filled = terete, open = broadleaved). **Significance codes: ‘**’ P < 0.01, ‘*’ P < 0.05.**

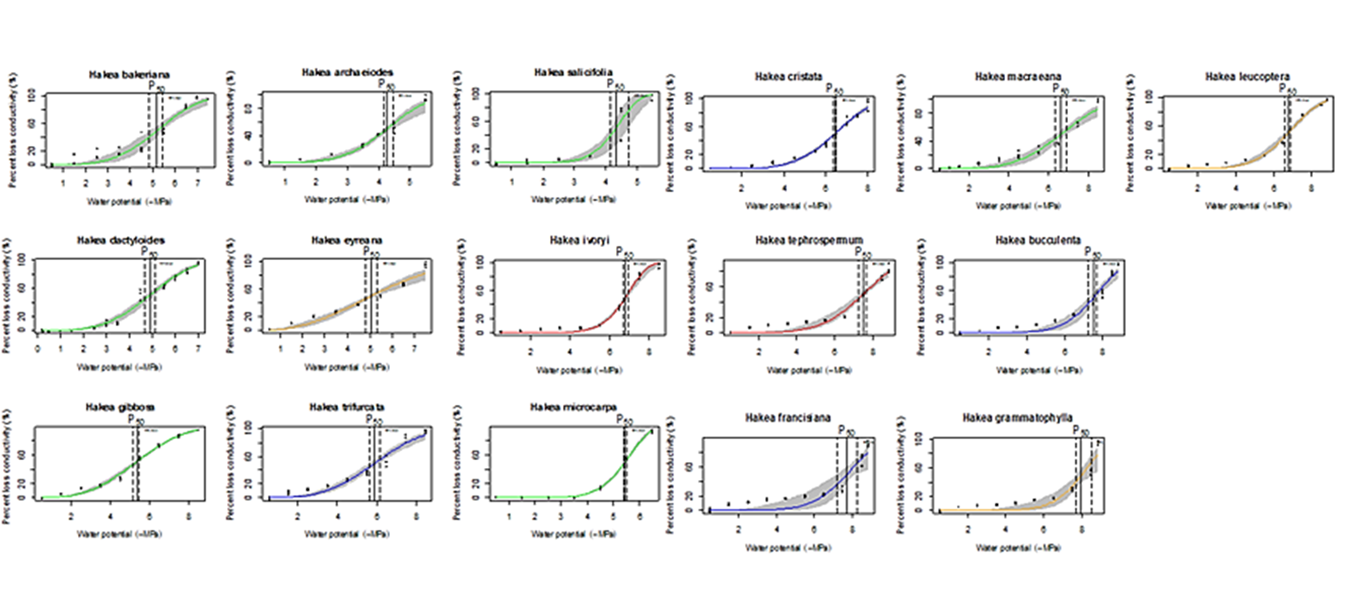

Fig S5: Hydraulic vulnerability curves for *Hakea* species showing the percentage loss of conductance against the stem water potential measured from the Cavitron. The curve was derived from combine measured of 2-3 branches for each species. Colour line represent vegetation type; green colour line = forest, blue = mediterranean, red = grassland and orange = arid.
